# The Wild Worm Codon Adapter: a web tool for automated codon adaptation of transgenes for expression in non-*Caenorhabditis* nematodes

**DOI:** 10.1101/2021.02.18.431865

**Authors:** Astra S. Bryant, Elissa A. Hallem

## Abstract

Advances in genomics techniques are expanding the range of nematode species that are amenable to transgenesis. Due to divergent codon usage biases across species, codon optimization is often a critical step for the successful expression of exogenous transgenes in nematodes. Platforms for generating DNA sequences codon optimized for the free-living model nematode *Caenorhabditis elegans* are broadly available. However, until now such tools did not exist for non-*Caenorhabditis* nematodes. We therefore developed the Wild Worm Codon Adapter, a tool for rapid transgene codon optimization for expression in non-*Caenorhabditis* nematodes. The app includes built-in optimization for parasitic nematodes in the *Strongyloides, Nippostrongylus* and *Brugia* genera as well as the predatory nematode *Pristionchus pacificus*. The app also supports custom optimization for any species using user-provided optimization rules. In addition, the app supports automated insertion of synthetic or native introns, as well as the analysis of codon bias in transgene and native sequences. Here, we describe this web-based tool and demonstrate how it may be used to analyze genome-wide codon bias in *Strongyloides* species.

## INTRODUCTION

Parasitic nematodes, including soil-transmitted gastrointestinal parasites in the genus *Strongyloides* and filarial nematodes such as *Brugia malayi,* are a major source of disease and economic burden (Lustigman *et al.* 2012). While *C. elegans* is often used as a model system for the study of parasitic nematodes, parasitic nematodes are behaviorally and genetically divergent from *C. elegans*; for example, they engage in a number of parasite-specific behaviors such as host seeking, host invasion, and intra-host migration (Haas 2003; Gang and Hallem 2016). Establishing methods that allow researchers to genetically manipulate parasitic nematodes directly is critical for understanding the genetic and cellular basis of parasitism in these species.

Historically, the application of functional genomics techniques in parasitic nematodes has lagged behind their use in the free-living model nematode *Caenorhabditis elegans*, due to multiple factors including the limited availability of genomic information (Castelletto *et al.* 2020). Recently released high-quality reference genomes for many parasitic nematode species (Hunt *et al.* 2016; Howe *et al.* 2017; International Helminth Genomes Consortium 2019) are a critical resource for parasitic nematode functional genomics techniques such as transgenesis and CRISRP/Cas9-mediated mutagenesis (Lok *et al.* 2017; Castelletto *et al.* 2020; Liu *et al.* 2020). The generation of transgenic parasitic nematodes is an essential tool for mechanistic studies of parasite-specific behaviors (Bryant *et al.* 2018; Gang *et al.* 2020). As our technical understanding of nematode genomics continues to develop beyond *C. elegans*, establishing accessible tools that automate the transgene design process for a broad selection of nematode species will greatly facilitate the application of genomics techniques in these species.

Transgenesis protocols are increasingly well-established in non-*Caenorhabditis* nematode species, including three soil-transmitted gastrointestinal parasites in the *Strongyloididae* family – the human parasite *Strongyloides stercoralis*, the rodent parasite *Strongyloides ratti*, and the Australian brushtail possum parasite *Parastrongyloides trichosuri* – as well as the rodent gastrointestinal parasite *Nippostrongylus brasiliensis*, the human-parasitic filarial nematode *Brugia malayi,* the predatory nematode *Pristionchus pacificus*, and the free-living nematodes *Auanema rhodensis* and *Auanema freiburgensis* (Grant *et al.* 2006; Lok *et al.* 2017; Adams *et al.* 2019; Castelletto *et al.* 2020; Han *et al.* 2020). Intragonadal microinjection, in which exogenous DNA is injected into either the syncytial gonad of free-living adult females (*Strongyloididae* species, *P. pacificus*), the distal gonad of free-living adult females (*Auanema* species), or the uterus of parasitic adult females (*B. malayi*), has been used to generate progeny expressing a range of transgenes (Schlager *et al.* 2009; Lok *et al.* 2017; Hong *et al.* 2019; Adams *et al.* 2019; Castelletto *et al.* 2020; Carstensen *et al.* 2020; Han *et al.* 2020). Intragonadal microinjection has also been used to achieve CRISPR/Cas9-mediated mutagenesis in *S. stercoralis*, *S. ratti*, *P. pacificus*, and *Auanema* species (Witte *et al.* 2015; Lok *et al.* 2017; Gang *et al.* 2017; Bryant *et al.* 2018; Adams *et al.* 2019; Han *et al.* 2020). In *B. malayi*, transfection of infective larvae has been used to deliver reporter plasmids and CRISPR constructs (Liu *et al.* 2018, 2020). Most recently, lentiviral transduction of infective larvae was used to deliver RNA interference molecules and drive expression of fluorescent reporters in *N. brasiliensis* (Hagen *et al.* 2021).

In non-*Caenorhabditis* nematodes and other species, successful transgene expression often requires the use of species-specific preferred codons. Codon usage bias is pervasive in species from all taxa, including nematodes: most amino acids may be encoded by multiple synonymous codons, and individual species tend to favor a specific set of codons, particularly for encoding highly expressed genes (Sharp and Li 1987; Mitreva *et al.* 2006; Cutter *et al.* 2006). The use of preferred codons is thought to promote efficient translation, although the exact mechanisms are not clear (Plotkin and Kudla 2011). Codon usage bias is believed to regulate the expression of exogenous transgenes as well as endogenous genes (Redemann *et al.* 2011). In non-*Caenorhabditis* nematodes, expression of exogeneous genes from transgenes is promoted by the use of species-specific codon usage patterns (Han *et al.* 2020; Hagen *et al.* 2021).

Although the process of codon-adapting transgenes for *C. elegans* is simplified by web-based platforms (Grote *et al.* 2005; Redemann *et al.* 2011), transgenes codon-adapted for other nematode species were previously designed by hand. Therefore, we created a web-based application, the Wild Worm Codon Adapter, which automates the process of codon optimization for transgenic expression in non-*Caenorhabditis* nematode species. Furthermore, the application permits automated insertion of synthetic or native introns into codon-optimized cDNA sequences, inclusion of which can significantly increase gene expression (Junio *et al.* 2008; Li *et al.* 2011; Crane *et al.* 2019; Han *et al.* 2020). Finally, the app enables users to rapidly assess relative codon bias of transgene sequences, as well as native *Strongyloides* and *C. elegans* genes, using genus-specific codon adaptation indices.

## MATERIALS AND METHODS

### Data source and preferred codon selection

Codon usage rules for *Strongyloides*, *Brugia*, *Nippostrongylus,* and *C. elegans* were calculated based on previously published codon count data (File S1). For *Strongyloides*, count data are from 11,458 codons from the 50 most abundant *S. ratti* expressed sequence tag (EST) sequences (Mitreva *et al.* 2006). For *C. elegans*, data are from 178 genes (73,164 codons) with the highest bias toward translationally optimal codons (Sharp and Bradnam 1997). For *Brugia* and *Nippostrongylus*, count data from 6,483 *B. malayi* sequences (561,296 codons) and 650 *N. brasiliensis* sequences (75,934 codons) were retrieved from Nematode.net v.4.0 (Martin *et al.* 2015).

Count data was used to quantify the relative adaptiveness of individual codons: the frequency that codon “i” encodes amino acid “AA” ÷ the frequency of the codon most often used for encoding amino acid “AA” (Sharp and Li 1987; Jansen *et al.* 2003). Preferred codons were defined as the codon with the highest relative adaptiveness value for each amino acid. Preferred codons for *Pristionchus* species (File S1) are based on the codon usage bias of the top 10% of expressed genes (Han *et al.* 2020).

### Codon adaptation index

The codon biases of individual sequences are quantified by calculating a Codon Adaptation Index (CAI), defined as the geometric average of relative adaptiveness of all codons in the sequence (Sharp and Li 1987; Jansen *et al.* 2003). CAI values relative to species-specific relative adaptiveness values are calculated using the seqinr package v3.6-1. CAI values are not calculated relative to *Pristionchus* or user-provided optimization rules, since in these cases optimal codons are defined directly rather than after calculating relative adaptiveness from count data.

### Fractional G-C content

The fraction of G+C bases in a sequence is calculated using the seqinr package v3.6-1.

### Intron insertion

Intron sequences are either: the three canonical artificial intron sequences used for *C. elegans* (Xu *et al.* 1996), *P. pacificus* native introns (Han *et al.* 2020), or Periodic A_n_/T_n_ Cluster (PATC)-rich introns from the *C. elegans* gene *smu-2* (introns 3-5) (Aljohani *et al.* 2020). For intron placement, the optimized cDNA sequence is divided at three predicted intron insertion sites spaced approximately equidistantly. Insertion sites are predicted *C. elegans* exon splice sites: preferably the stringent consensus sequences (5’-AAG^G-3’, 5’-AAG^A-3’, 5’-CAG^G-3’, 5’-CAG^A-3’), however minimal consensus sequences (5’-AG^G-3’, 5’-AG^A-3’) are used if more stringent sites are not present or if there are fewer insertion sites than the requested number of introns (Meyer *et al.* 1997; Redemann *et al.* 2011). The user-specified number of introns (up to a maximum of three) are inserted into the sequence using the 5’ insertion site first and continuing in the 3’ direction (Crane *et al.* 2019; Han *et al.* 2020).

### cDNA sequence lookup

In “Analyze Sequences” mode, users may submit search terms including: stable gene IDs, *C. elegans* gene names, or matched gene IDs and cDNA sequences as either a two-column .csv file or an .fa file. If users supply only gene IDs, either via a text box or file upload, the app first fetches the cDNA sequences from WormBase ParaSite via the biomaRt package v2.42.1 (Durinck *et al.* 2005, 2009). The following types of gene IDs may be used: stable gene or transcript IDs with prefixes “SSTP,” “SRAE,” “SPAL,” “SVE,” or “WB;” *C. elegans* stable transcript IDs; or *C. elegans* gene names prefaced with the string ‘Ce-’ (*e.g., Ce-tax-4*).

### Genome-wide codon bias and Gene Ontology (GO) analysis

FASTA files containing all coding sequences (CDS) for *S. stercoralis*, *S. ratti*, *S. papillosus*, *S. venezuelensis*, *N. brasiliensis*, *B. malayi*, and *C. elegans* were downloaded from WormBase ParaSite (WBPS15) and analyzed using the app. For *C. elegans* and *Strongyloides* species, results were filtered to identify six functional subsets: the 2% of genes with highest and lowest *S. ratti* CAI values, the 2% of genes with the highest and lowest *C. elegans* CAI values, and the 2% of genes with highest and lowest RNA-seq expression in free-living females. Log_2_ counts per million (CPM) expression in free-living adult females was downloaded from the *Strongyloides* RNA-seq Browser (Bryant *et al.* 2021). For statistical comparisons of the expression of the highest and lowest *Strongyloides*-codon-adapted genes, relative to all genes, a 2-way ANOVA (type III) with Tukey *post-hoc* tests was performed in R using the car package v3.0-8. GO analyses of functional subsets were performed using the gprofiler2 package v0.1.9, with a false discovery rate (FDR)-corrected *p*-value < 0.05. Commonly enriched GO terms in each subset were defined as GO terms that were enriched in all four of the *Strongyloides* species (or at least three *Strongyloides* species in the case of gene-expression-based subsets) with an FDR-corrected *p*-value of ≤ 0.001. For statistical comparisons of genome-wide CAI values and fractional G-C content between species, Kruskal-Wallis tests with *post-hoc* Dunn’s tests were performed in R using the dunn.test package v1.3.5. For *post-hoc* Dunn’s tests, *p*-values were corrected using the Bonferroni method.

### Data Availability

Preprocessing and analysis source code, plus codon frequency data, intron sequences, and supplemental files are available at: https://github.com/HallemLab/Bryant-and-Hallem-2021

A web-hosted version of the app is available at: https://hallemlab.shinyapps.io/Wild_Worm_Codon_Adapter/

App source code and deployment instructions are available at: https://github.com/HallemLab/Wild_Worm_Codon_Adapter

The following supplemental files have been uploaded to figshare. File S1 contains codon usage frequencies and optimal codons for: highly abundant *S. ratti* expressed sequence tag (EST) transcripts (Mitreva *et al* 2006), highly expressed *C. elegans* genes (Sharp and Bradnam 1997), highly expressed *P. pacificus* genes (Han *et al* 2020), *B. malayi* contig sequences (http://www.nematode.net) (Martin *et al.* 2015), *N. brasiliensis* contig sequences (http://www.nematode.net) (Martin *et al.* 2015), and all *S. ratti* ESTs (Mitreva *et al* 2006). File S2 contains a code freeze for the Wild Worm Codon Adapter. File S3 contains gene IDs, CAI values, GC ratios, and GO term accession numbers of the 2% of genes with the highest and lowest CAI values for each *Strongyloides* species and *C. elegans*. File S4 contains GO analysis results for the 2% of genes with the highest and lowest CAI values for each *Strongyloides* species and *C. elegans*. File S5 contains GO terms significantly enriched in all four *Strongyloides* species for the highest (top 2%) and lowest (bottom 2%) *Strongyloides* codon-adapted sequences, as well as GO terms significantly enriched in at least three *Strongyloides* species for genes with the highest (top 2%) and lowest (bottom 2%) expression in free-living females.

## RESULTS AND DISCUSSION

### Software Functionality

The Wild Worm Codon Adapter app features two usage modes: “Optimize Sequences” mode and “Analyze Sequences” mode (Figures 1–2, File S2). “Optimize Sequences” mode automates the process of transgene codon adaptation and intron insertion. For codon optimization, the app features built-in preferred codons for four non-*Caenorhabditis* nematode genera for which transgenesis protocols are increasingly well-established: *Strongyloides*, *Nippostrongylus*, *Pristionchus*, and *Brugia*. This mode also supports custom codon optimization based on a user-provided list of preferred codons.

**Figure 1.**
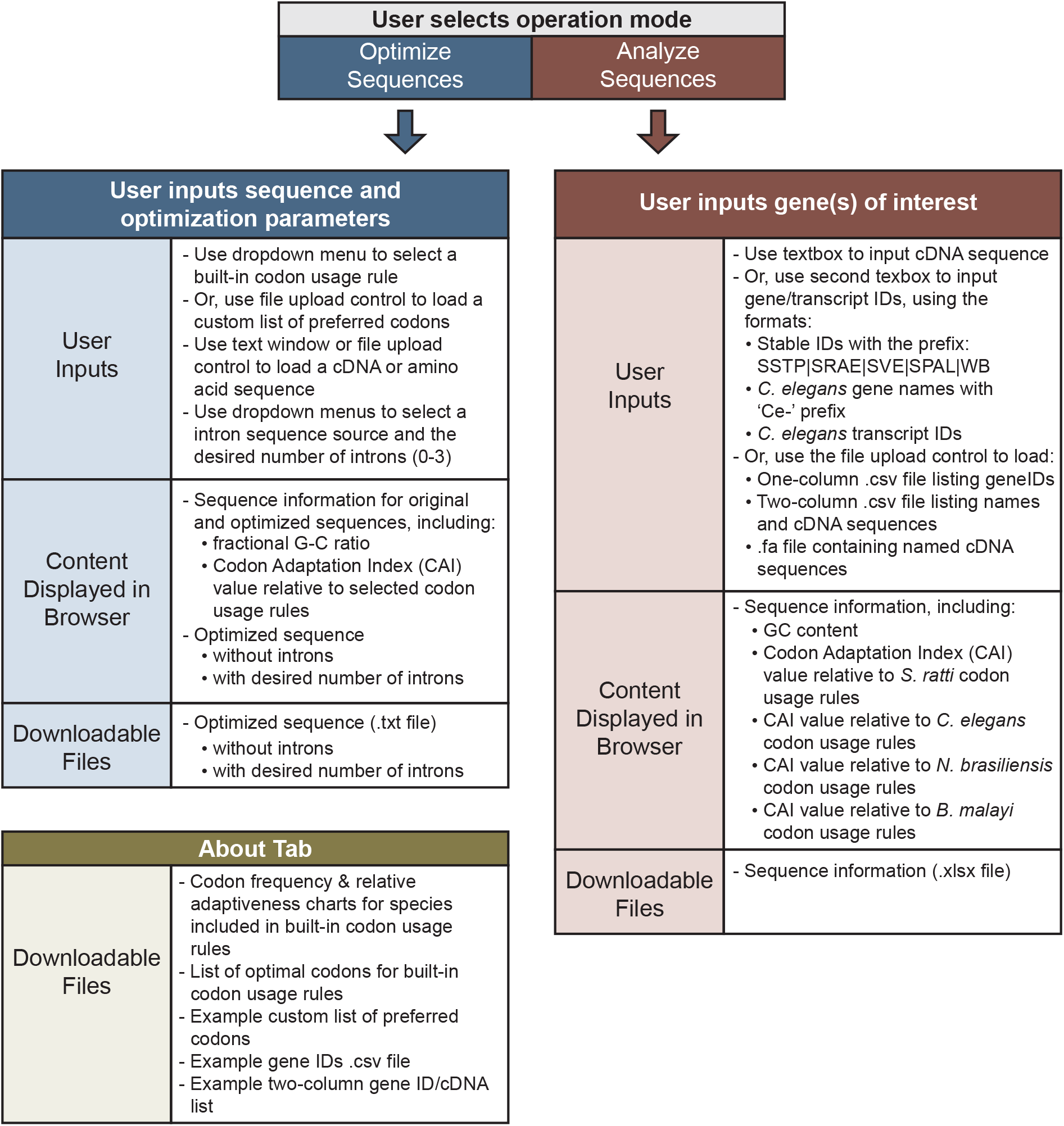
A UI overview of the Wild Worm Codon Adapter app. The overview shows user inputs, content displayed in the browser, and downloadable files. The application features two modes: Optimize Sequences (blue panels) and Analyze Sequences (red panels) that are accessed via separate tabs in the browser window. The app also includes an About tab that presents methods information and a menu for downloading files. Available files include example input files as well as tables of species-specific optimal codons, species-specific codon frequencies, and relative adaptiveness values used to calculate codon adaptation indices.

**Figure 2.**
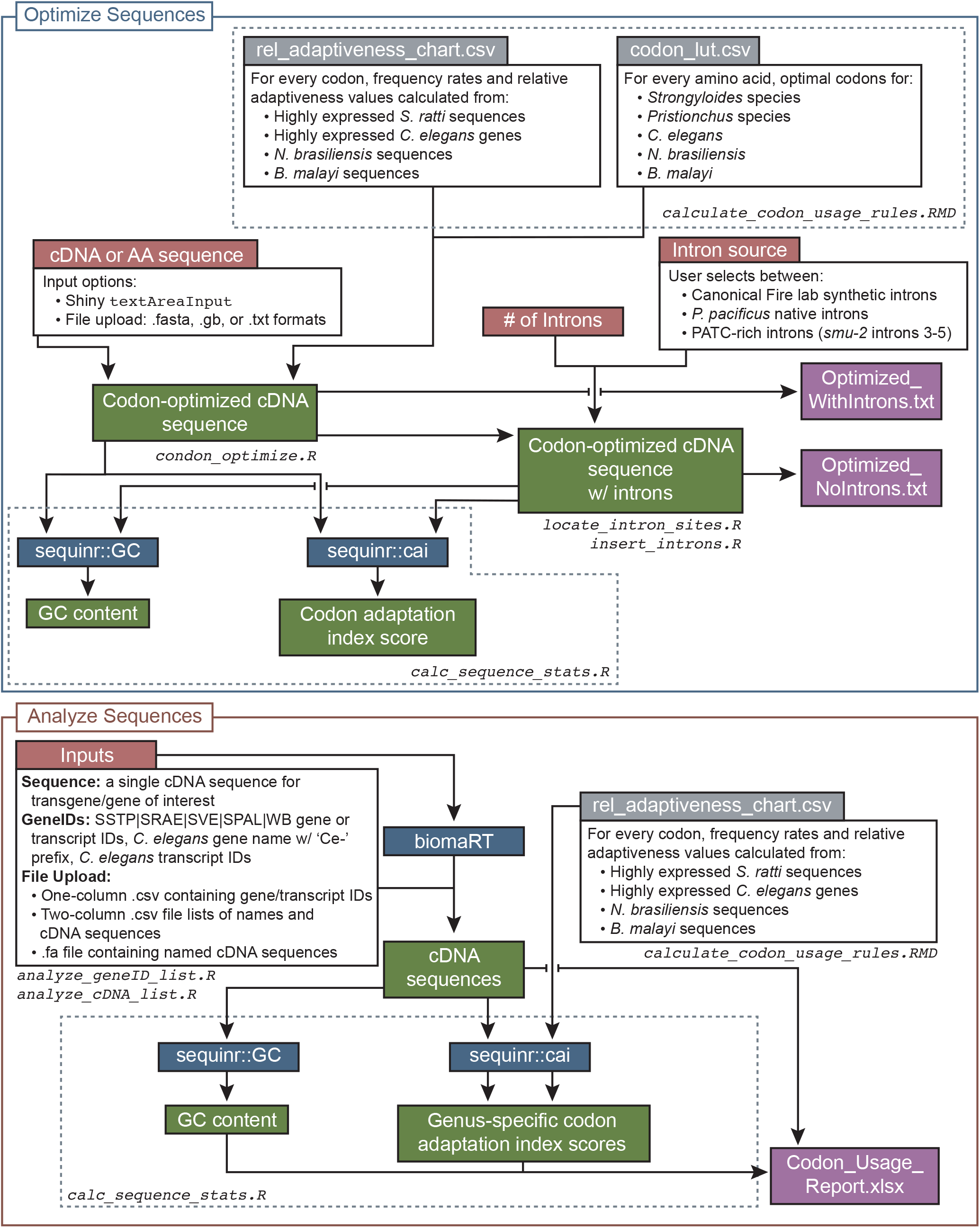
Detailed graphic view of the codebase of the Wild Worm Codon Adapter app. The application features two usage modes: Optimize Sequences (upper, blue panel) and Analyze Sequences (lower, red panel). Grey boxes are preprocessed data inputs generated from published data. Red boxes are user inputs. Green boxes are output elements displayed within the browser; output values directly depend on inputs provided via red elements. Blue boxes are commands run in R. White boxes are code details or input options. Purple boxes are downloadable output file options. Dashed lines show division of code elements into the named files.

To generate codon-optimized sequences suitable for expression in *Strongyloides, Nippostrongylus*, *Brugia,* or *Pristionchus* species, users select the appropriate codon usage rule and then submit cDNA or amino acid sequences via a text window or file upload. To codon-optimize transgenes for expression in any additional organism of interest, users may prefer to supply a custom list of preferred codons using the following format: a 2-column .csv file listing single-letter amino acid symbols and corresponding 3-letter optimal codon sequence; only one optimal codon should be provided per amino acid and stop codons should be designated using the ‘*’ symbol. An example custom preferred codon table is available to download from the website.

For *Strongyloides* and *Pristionchus* species, codon optimization is based on *S. ratti* and *P. pacificus* codon usage patterns (Mitreva *et al.* 2006; Han *et al.* 2020). Codon optimization for *Brugia* species is based on genome-wide codon bias in *B. malayi*. For *Nippostrongylus* species, optimization rules are based on *N. brasiliensis* codon usage biases. Codon usage is often well conserved between closely related nematode species (Mitreva *et al.* 2006; Han *et al.* 2020). Thus, codon usage rules generated from individual species are likely effective across closely related species (*e.g.*, across members of a genus).

We calculated built-in codon usage rules for *Strongyloides* and *Pristionchus* from the codon usage patterns observed in highly expressed genes (Mitreva *et al.* 2006; Han *et al.* 2020), since the codon usage patterns of highly expressed genes are thought to correlate with higher protein expression (Sharp and Li 1987; Plotkin and Kudla 2011). However, the codon usage rules for highly expressed *S. ratti* genes are extremely similar to those observed across all *S. ratti* coding sequences (File S1). In contrast, the preferred codon usage rules we implement for *Strongyloides*, *Pristionchus*, and *Brugia* are distinct from the *C. elegans* codon usage rules (File S1), consistent with observations that individual transgenes show limited expression across nematode species with divergent genomes (Hunt *et al.* 2016; Lok *et al.* 2017; Castelletto *et al.* 2020; Han *et al.* 2020).

Previous studies have suggested that highly divergent codon usage patterns between nematodes are driven in part by the extreme AT-richness of some nematode genomes (Mitreva *et al.* 2006; Cutter *et al.* 2006; Hunt *et al.* 2016; Han *et al.* 2020). As expected, in *S. ratti* and *B. malayi*, which have AT-rich genomes, the preferred codons that comprise the built-in usage rules are AT-biased (preferred codon fractional G-C content: *S. ratti* = 0.33, *B. malayi* = 0.32, *N. brasiliensis* = 0.52, *P. pacificus* = 0.46, *C. elegans* = 0.52). Thus, codon optimization for expression in *Strongyloides* and *Brugia* species will likely yield AT-rich optimized sequences. Given the use of a single codon sequence per amino acid and the AT-bias in preferred codons for some species, users may want to eliminate repeated nucleotide patterns, hairpin loops, or unwanted restriction sites prior to final gene synthesis.

To insert introns into optimized cDNA sequences, users select the type and number of introns for insertion into the optimized sequence (up to three). Users may choose between three sets of unique intron sequences: canonical *C. elegans* artificial introns (Xu *et al.* 1996), *P. pacificus* native introns (Han *et al.* 2020), or Periodic A_n_/T_n_ Cluster (PATC)-rich introns that enhance germline expression of transgenes in *C. elegans* (Aljohani *et al.* 2020). The app identifies three putative intron insertion sites spaced approximately equidistantly within the optimized cDNA sequence, and inserts the user-specified type and number of introns (Xu *et al.* 1996; Meyer *et al.* 1997; Redemann *et al.* 2011). In *C. elegans* and *P. pacificus*, a single 5’ intron is sufficient for intron-mediated enhancement of gene expression, whereas a single 3’ intron is not (Crane *et al.* 2019; Han *et al.* 2020). Thus, when the user chooses to insert fewer than three introns, the three hypothetical insertion sites are filled as needed, starting from the 5’ site.

The app displays the optimized sequence with and without introns; users may download these sequences as a .txt files. “Optimize Sequences” mode reports the fractional G-C content for the original and optimized sequences. When optimizing using *Strongyloides*, *Nippostrongylus*, or *Brugia* usage rules, the app also provides a measure of the codon usage bias for both the original and optimized cDNA sequences by reporting codon adaptation index (CAI) values relative to the selected usage rule. When optimizing using *Pristionchus* or user-provided optimization rules, since optimal codons are defined directly, the app does not calculate CAI values.

Finally, the “Analyze Sequences” mode (Figure 1) was designed to support descriptive analyses of transgene sequences as well as native *Strongyloides* or *C. elegans* coding sequences. To measure how well codon-adapted a transgene is for expression in *Strongyloides*, *Brugia*, *Nippostrongylus*, or *C. elegans*, users may submit the relevant cDNA sequence(s). For individual cDNA sequences, the following values are calculated and displayed as a downloadable table: fractional G-C content; and codon adaptation index values relative to *Strongyloides*, *Brugia*, *Nippostrongylus*, and *C. elegans* codon usage rules (*Sr*-CAI, *Bm*-CAI, *Nb*-CAI, and *Ce*-CAI, respectively). When presented with a list of *Strongyloides* or *C. elegans* gene or transcript IDs, the app first retrieves associated cDNA sequences from WormBase ParaSite before calculating the quantifications listed above.

### Benchmarking and Example Usage

To benchmark the use of the codon adaptation index to quantify codon adaptiveness of native genes, we used “Analyze Sequences” mode to assess and compare genome-wide codon bias patterns in *Strongyloides* species, *B. malayi*, *N. brasiliensis*, and *C. elegans*. *P. pacificus* was not included because the app does not calculate CAI values relative to *P. pacificus* codon usage rules. Consistent with previous findings (Mitreva *et al.* 2006), we found that the distribution of genome-wide codon bias varied by genus, such that the genus-specific CAI values of *C. elegans* coding sequences were lower than those for parasitic species (Figure 3A; *p* < 0.0001 for all comparisons to *C. elegans*, Kruskal-Wallis test with Dunn’s *post-hoc* tests). Also consistent with previous observations that *Strongyloides* genomes are highly AT-rich (Mitreva *et al.* 2006; Cutter *et al.* 2006; Hunt *et al.* 2016), we observed that *Strongyloides* coding sequences displayed lower fractional G-C content than *C. elegans*, *N. brasiliensis*, and *B. malayi* coding sequences (Figure 3B; *p* < 0.0001 for *Strongyloides* species grouped together and compared to each remaining species, Kruskal-Wallis test with Dunn’s *post-hoc* tests). Together these observations emphasize the likely benefit of transgene codon-optimization for promoting successful transgenesis in non-*Caenorhabditis* species. Finally, consistent with our use of *Strongyloides* codon usage rules based on highly abundant *S. ratti* ESTs, the highest and lowest *Strongyloides*-codon-adapted sequences generally displayed significantly higher and lower expression, respectively, in free-living female life stages relative to the expression of all genes in the genome (Figure 3C). However, the range of gene expression values between the highest and lowest codon-adapted sequences are largely overlapping; thus, the degree of codon adaptation exhibited by individual genes is not the sole determinant of mRNA expression level in these species.

**Figure 3.**
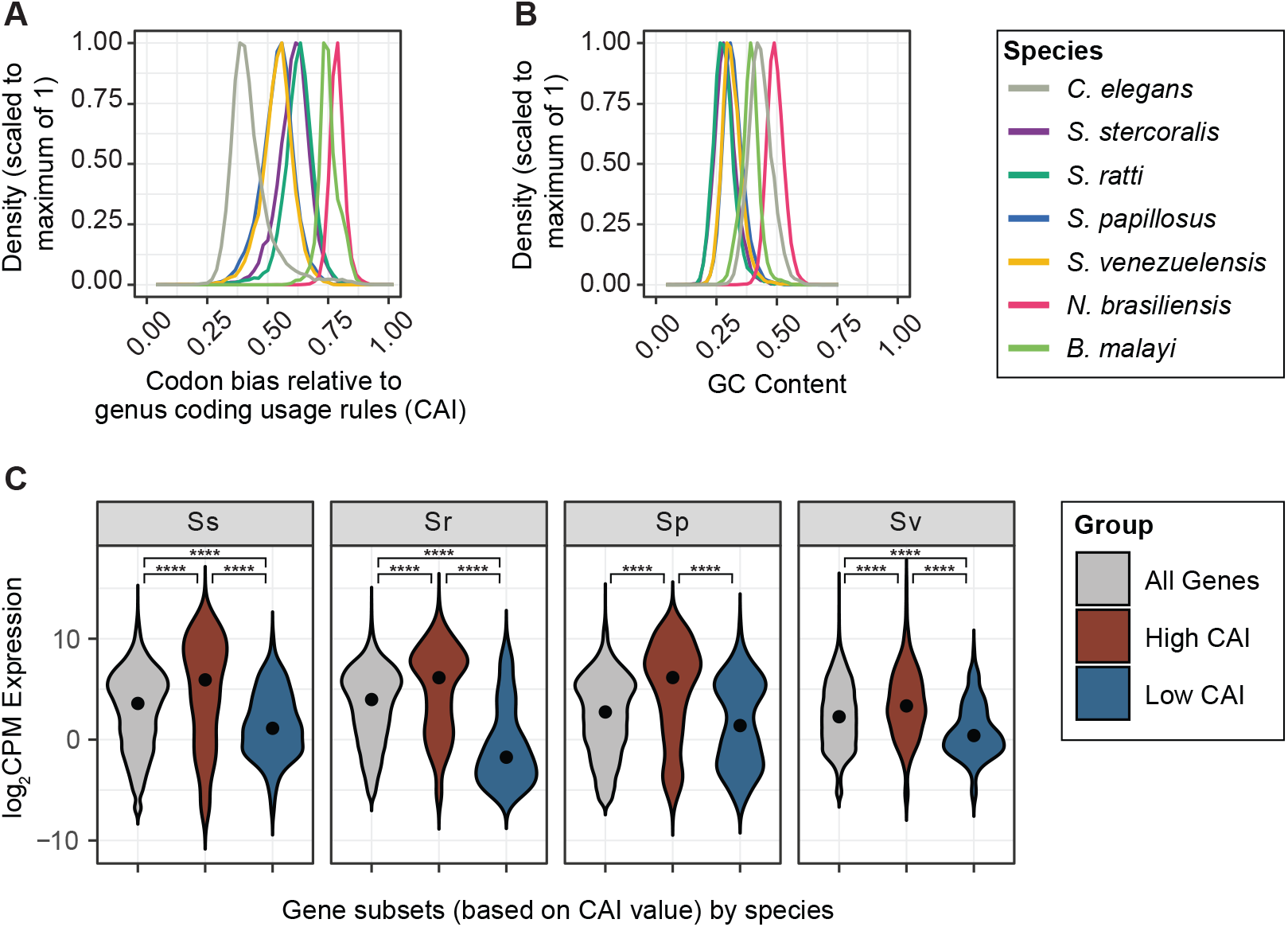
Codon adaptiveness and GC content across species. **A)** Density plot of codon adaptation index (CAI) values in a species’ genome. Codon adaptiveness varies between species; *C. elegans* genome-wide codon adaptiveness is significantly lower than other species (*p* < 0.0001 comparing *C. elegans* to each remaining species, Kruskal-Wallis test with Dunn’s *post-hoc* tests). For each species, codon bias is calculated relative to the genus-level codon usage rules (File S1). **B)** Density plot of fractional GC content across species. Consistent with previous reports (Mitreva *et al.* 2006; Cutter *et al.* 2006; Hunt *et al.* 2016), *Strongyloides* coding sequences tend to be more AT-rich than the coding sequences of other species (*p* < 0.0001 for the grouped *Strongyloides* species versus each remaining species, Kruskal-Wallis test with Dunn’s *post-hoc* tests). For both panels, values were calculated by submitting a list of all predicted coding sequences to the Wild Worm Codon Adapter app in Analyze Sequences mode. **C)** Violin plot of log_2_ counts per million (CPM) expression in free-living females for all genes (grey), genes in the top 2% of CAI values for each species (red), and genes in the bottom 2% of CAI values for each species (blue). Dots indicate median values. \*\*\*\**p* < 0.00005, 2-way ANOVA with Tukey *post-hoc* tests.

Next, we used gene ontology (GO) analyses to assess the putative functions of the most highly or poorly *Strongyloides-*codon adapted coding sequences (defined as the top/bottom 2% of *Sr*-CAI values; Files S3, S4). For the highest *Strongyloides*-codon-adapted sequences, GO terms that are significantly enriched in all four *Strongyloides* species are primarily associated with structural integrity and ribosomal components (Figure 4A-B, File S5). As expected, given that the *Strongyloides* codon usage rules are based on highly abundant sequences, there is significant overlap between GO terms enriched in the top 2% of *Strongyloides*-codon-adapted sequences and GO terms enriched in the genes that are most highly expressed in free-living females (Figure 4B, File S5). Despite the differences between codon usage patterns in *Strongyloides* and *C. elegans,* GO terms enriched in the top 2% of *Strongyloides*-codon-adapted sequences are also significantly enriched in the top 2% of *C. elegans*-codon-adapted sequences (*i.e., C. elegans* sequences that display high codon usage bias relative to *C. elegans* codon usage rules) (Figure 4B). In contrast, for parasite sequences that are the least *Strongyloides-*codon-adapted, commonly enriched GO terms are associated with DNA metabolism and interactions (Figure 4C-D, File S5); these terms are not enriched in low-expressing *Strongyloides* genes or poorly *C. elegans-*codon-adapted *C. elegans* sequences (Figure 4D, File S5). Although the causes and consequences of codon bias are not fully understood (Sharp *et al.* 2010; Plotkin and Kudla 2011), our observations of a common set of GO terms enriched in both *Strongyloides-* and *C. elegans*-codon-adapted genes suggest that in nematodes, heighted codon bias reflects a non-random process that systematically imposes divergent codon usage patterns on genes associated with a common set of biological functions.

**Figure 4.**
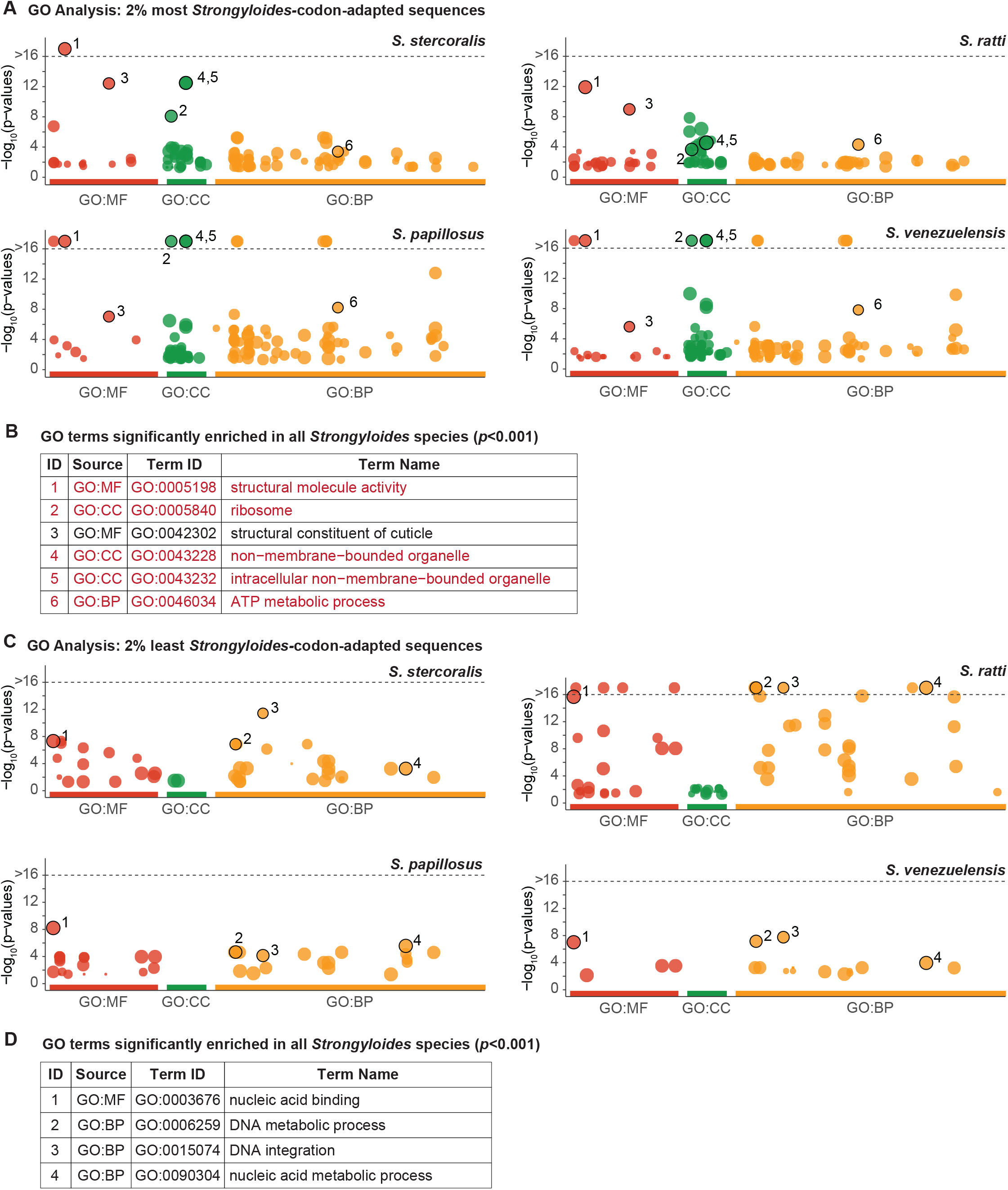
Functional enrichment of highly and poorly codon-adapted genes. Gene Ontology (GO) enrichment across *Strongyloides* species for sequences displaying high and low degrees of *Strongyloides* codon bias. **A-B)** Manhattan plots and table showing GO enrichment for sequences in the top 2% of CAI values for each species. X-axis labels (A) and table Source values (B) indicate the three GO vocabulary categories: biological process (BP), molecular function (MP) and cellular component (CC). Numbered circles (A) and table ID numbers (B) indicate GO terms that are significantly enriched (*p* ≤ 0.001) in all four *Strongyloides* species. For Manhattan plots, −log_10_(*p*-values) along the y-axis are capped if ≥ 16; this threshold is indicated by a grey dashed line. All GO terms included in panel B are also significantly enriched in the top 2% of *C. elegans*-codon-adapted *C. elegans* sequences. Red terms are also significantly enriched among genes with the highest expression (top 2%) in free-living females, in at least 3 *Strongyloides* species. **C-D)** Manhattan plots and table of GO enrichment for sequences that compose the bottom 2% of CAI values. None of the GO terms included in panel D are also significantly enriched (*p* ≤ 0.001) in poorly *C. elegans*-codon-adapted *C. elegans* sequences or in genes with the lowest expression (bottom 2%) in free-living females. For all plots, *p-*values are false discovery rate (FDR)-corrected.

## CONCLUSIONS AND FUTURE DIRECTIONS

Codon optimization of transgenes significantly improves the likelihood of successful transgenesis in non-*Caenorhabditis* nematodes. Here, we present the Wild Worm Codon Adapter, a web-based tool designed to replicate and extend the functionality of popular codon-optimization tools to include non-*Caenorhabditis* nematode genera such as *Strongyloides*, *Pristionchus, Nippostrongylus*, and *Brugia* (Grote *et al.* 2005; Redemann *et al.* 2011). The open-source code is extendable; users may perform codon-optimization for an unlimited selection of species by providing a custom list of optimal codons, and additions to the list of built-in optimization rules can be implemented as requested. Going forward, we hope that the Wild Worm Codon Adapter will simplify the process of transgene design for researchers using functional genomics to study non-*Caenorhabditis* nematodes.

## ACKNOWLEDGMENTS

The authors would like to thank Dr. Stephanie DeMarco and Dr. Michelle Castelletto for helpful discussion, and Dr. Ruhi Patel for helpful comments on the manuscript.

